# Electrical Impedance Spectroscopy as a Tool to Detect the Epithelial to Mesenchymal Transition in Prostate Cancer Cells

**DOI:** 10.1101/2024.09.29.615724

**Authors:** Lexi L. Crowell, Luis A. Henriquez, Mary Tran, Tayloria N.G. Adams

## Abstract

Prostate cancer (PCa) remains a significant health threat, with chemoresistance and recurrence posing major challenges despite advances in treatment. The epithelial to mesenchymal transition (EMT), a biochemical process where cells lose epithelial features and gain mesenchymal traits, is linked to chemoresistance and metastasis. Electrical impedance spectroscopy (EIS), a novel label-free electrokinetic technique, offers promise in detecting cell phenotype changes. In this study, we employed EIS to detect EMT in prostate cancer cells (PCCs). PC3, DU145, and LNCaP cells were treated with an EMT induction media for five days. EIS characterization revealed unique impedance spectra correlating with metastatic potential, distinguishing DU145 EMT+ and EMT-cells, and LNCaP EMT+ and EMT-cells (in combination with dielectrophoresis), with comparisons made to epithelial and mesenchymal controls. These changes were supported by shifts in electrical signatures, morphological, and protein expression, including downregulation of E-cadherin and upregulation of vimentin. No phenotype change was observed in PC3 cells, which maintained a mesenchymal phenotype. EMT+ cells were also distinguishable from mixtures of EMT+ and EMT-cells. This study demonstrates key advancements: application of EIS and dielectrophoresis for label-free EMT detection in PCCs, characterization of cell electrical signature after EMT, and EIS sensitivity to EMT transitions. Detecting EMT in PCa is important to the development of more effective treatments and overcoming the challenges of chemoresistance.

## 1. Introduction

In the United States, prostate cancer (PCa) poses a significant health threat and has the highest incidence of cancer amongst men, with an estimated 300,000 new cases and 35,000 deaths expected in 2024. Yet, significant gains have been made in survival rates, with a 50% decrease in deaths since the mid-1990s [1]. With advances in medicine, patients may undergo surgical, radiotherapy, and chemotherapy treatments to combat PCa [2]. However, even after radical prostatectomy, there remains a possibility for cancer recurrence [3], representing a progression in PCa that is challenging to treat. The probability of recurrence lies between 15 and 40% in a span of 10 years [3]. Recurrence may occur locally, or the cancer may metastasize, leading to death. Consequently, addressing recurrence in a clinical setting is crucial for improving long-term patient outcomes.

PCa metastasis remains a challenge not only due to the spread of cancer to distant organs but also due to subsequent tumor heterogeneity and plasticity [4]. These factors impact the effectiveness of PCa treatments. One hallmark of metastasis is the epithelial to mesenchymal transition (EMT), a reversible process in which polarized epithelial cells undergo biochemical changes ultimately leading to transformation into mesenchymal cells [5], which is distinguishable by increased chemoresistance [6]. During EMT, cells upregulate the expression of mesenchymal markers such as N-cadherin and vimentin, leading to aggressiveness and invasiveness, while downregulating epithelial markers such as E-cadherin and ZO-1, resulting in the loosening of cell junctions, disruption of cell-to-cell contact, and the development of elongated, motile spindle-shaped cells with increase resistance to apoptosis [5, 7]. Subsequent effects of EMT include cells’ ability to self-renew and increases the heterogeneity of the cell population [8]. Several cell signaling pathways induce EMT including transforming growth factor beta (TGF-β), Wnt, notch, sonic hedgehog, and interleukin-6 [7]. Thus, cancer cells exist in two states: an epithelial state, characterized by high polarity, strong cell-to-cell adhesion, cobblestone morphology, and low chemoresistance, and a mesenchymal state, characterized by low polarity, reduced cellto-cell adhesion, spindle-like elongated morphology, high migratory capabilities, and increased chemoresistance [7].

Despite extensive research and the recognized role of EMT in chemoresistance, no approved clinical application for EMT markers currently exists. Consequently, alternative methods such as electrical impedance spectroscopy (EIS) are worthy of exploration due to their ability to detect phenotypic changes in cells [9]. EIS is a non-invasive electrokinetic technique that leverages the inherent electrical properties of cells to distinguish various types of cancer cells [10] and assess their dynamics processes, including adhesion [11], proliferation [12], spreading [13], motility [14], and apoptosis [15], as well as chemoresistance [16]. Briefly, EIS uses non-uniform electric fields to interact with cell membranes, membrane proteins, and intracellular molecules [17]. The resulting impedance spectra has frequency-dependent characteristics unique to the cell population being examined. For a comprehensive review of EIS theory, refer to [18] and [16]. At radio frequencies, the impedance spectra provides valuable insight into biological cells including, surface charge mobility, cell membrane thickness, and relaxation effects caused by proteins, amino acid residues, and internal organelles [17]. Thus, for prostate cancer cells (PCCs) selectively investigating radio frequency EIS responses can reveal intrinsic and extrinsic factors that contribute to cell phenotype such as those associated with EMT.

To the authors’ knowledge there are no studies using EIS to study EMT in PCCs. However, EIS has been used to distinguish PCC lines [9, 19], and other impedimetric techniques such as electric cell-substrate impedance sensing (ECIS) [20] and impedance flow cytometry (IFC) [21] have been used to monitor EMT in lung and breast cancer cells. If we expand to include electrochemical immunosensors than several studies have investigated EMT for liver, breast, ovarian, and pancreatic cancer cells [22, 23]. While all of these impedimetric techniques are valuable for studying EMT in cancer cells, our specific focus remains on EIS.

In this study, the EIS of three PCC lines that underwent EMT induction were characterized. The PC3 cell line, derived from a vertebral metastasis lesion of grade IV prostatic adenocarcinoma in a 62-year-old male [24], and the DU145 cell line, from a 69-year-old male with lymphocytic leukemia and advanced prostate carcinoma [25], were analyzed alongside the androgen-dependent LNCaP cell line, which originated from a metastatic lesion of prostatic adenocarcinoma in a male [26]. PCCs were treated with an EMT induction media over five days and characterized using EIS in a simple parallel electrode microfluidic device and, in some cases, by dielectrophoresis, an alternative electrokinetic technique. Our results showed that PC3, DU145, and LNCaP cells have unique electrical signatures without and with EMT treatment (EMT-and EMT+). We verified EMT induction with several outputs: discernible difference in the impedance spectra, comparison of the impedance spectra of PC3 and DU145 cells to epithelial (human primary prostate cells, HPrECs) and mesenchymal (human mesenchymal stem cells, hMSCs) controls, cell morphology assessment, and immunofluorescent imaging of E-cadherin and vimentin protein expression. Additionally, the dielectrophoresis analysis distinguished the LNCaP EMT+ and EMT-cells. These outputs confirmed that the EMT was successfully induced in the DU145 and the LNCaP cells but not in the PC3 cells. These findings demonstrate the potential of EIS to detect EMT-specific phenotype changes in PCCs and highlight several key advancements: the application of EIS and dielectrophoresis for label-free EMT detection in PCCs, characterization of changes in cell electrical signatures with EMT, and demonstrated of the sensitivity of EIS to EMT transitions.

## 2. Materials and Methods

### 2.1 Cell Culture

PC3, DU145, and LNCaP cells were obtained from American Type Culture Collection (ATCC, Manassas, VA, USA). These cells were subcultured in Roswell Park Memorial Institute (RPMI)-1640 medium (Thermo Fisher, A1049101) supplemented with 10% (v/v) heat-inactivated fetal bovine serum (Life Technologies, Carlsbad, CA, USA, 18121001), 50 U mL-1 penicillin and 50 µg mL-1 streptomycin (Life Technologies, Carlsbad, CA, 15140122) at 37°C in a humidified 5% CO_2_ incubator.

Umbilical cord-derived hMSCs obtained from ATCC (PCS-500-100) were subcultured in MSC basal medium (ATCC, PCS-500-030) supplemented with a growth kit, low serum (ATCC, PCS-500-040) at 37°C in a humidified 5% CO_2_ incubator. HPrECs were obtained from ATCC (PCS-440-010) and subcultured in prostate epithelial cell basal medium (ATCC, PCS-440-030) supplemented with prostate epithelial cell growth kit (ATCC, PCS-440-040) at 37°C in a humidified 5% CO_2_ incubator.

### 2.2 EMT Treatment

PC3, DU145, and LNCaP cells were treated with StemXVivio EMT-inducing media supplement (R&D Systems, Minneapolis, MN, USA, CM017) according to the manufacturer’s instructions. Specifically, two days after seeding 30,000 cells, the growth media was replaced with growth media supplemented with 1% EMT-inducing media. The EMT media was refreshed after 48 hours, and the cells were cultured in the EMT media for 5 days. On day 6, the cells were processed for electrokinetic, morphology, and immunofluorescent analyses.

### 2.3 Cell Preparation for Electrokinetic Characterization

After EMT treatment, the monolayer of cells was rinsed once with 1X Dulbecco’s phosphate-buffered saline (DPBS) (Life Technologies, Carlsbad, CA, USA) followed by trypsinization with 0.05% Trypsin-EDTA (Life Technologies, Carlsbad, CA, USA) for 5 minutes. Once cells were detached from the culture plate, they were neutralized with an equal volume of growth medium. The cell suspension was centrifuged at 134 x g for 5 minutes to pellet the cells. A low conductivity buffer (LCB) was made with Milli-Q water containing 8.5% (w/v) sucrose and 0.3% (w/v) D-glucose. The conductivity was adjusted to 100 ± 5 µS/cm using RPMI-1640 (Thermo Fisher, 11875135). The LCB was passed through a 22-micron filter for sterilization. Next, the cell pellet was washed three times in the LCB at 150 x g for 5 minutes. For the final resuspension the cell concentration was adjusted to 300,000 cells/mL. The following samples were assessed: PC3 EMT-, PC3 EMT+, DU145 EMT-, DU145 EMT+, LNCaP EMT-, LNCaP EMT+, as well as mixtures of PC3 EMT-/EMT+, DU145 EMT-/+, and LNCaP EMT-/+ cells.

### 2.4 Electrokinetic Characterization with EIS

For EIS, a simple microwell device with parallel electrodes was fabricated using previously published techniques [27, 28] with electrode dimensions of 50 µm width and 100 µm gap between the electrodes. The microfluidic device was washed 3 times with the LCB and attached to a Reference 600+ Potentiostat/Galvanostat/ZRA (Gamry Instruments, Warminster, PA, USA). The EIS characterization was completed using a frequency sweep from 350 Hz to 5 MHz at 10mV for 5 minutes. At least 3 to 5 technical replicates were completed for each independent experiment. This characterization was repeated for three biological replicates (n=3) of the PC3, DU145, and LNCaP cells. For hMSCs and HPrECs, the experiments were completed for three biological replicates (n=3).

### 2.5 Analysis of EIS Data

Once the EIS data was collected, outliers were removed using Prism 10. With the outliers removed, the technical replicates were normalized to eliminate any dependence on the LCB. Normalization was done by dividing the measured cell impedance at each frequency by the measured LCB impedance at the same frequency given by,

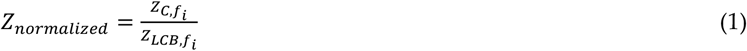

where *Z*_*normalized*_ is the normalized impedance, 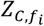 is the cell impedance at frequency *f*_i_, 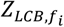 is the buffer impedance at frequency *f*_i_, and *i* represents each frequency in the sweep. The normalized impedance spectra for each of the biological replicates (n=3) of the PC3, DU145, and LNCaP cells were averaged to plot an average impedance spectrum. This process was repeated for the hMSCs and HPrECs. Additionally, the mean normalized impedance over the tested frequency range was calculated. A one-way ANOVA with Tukey’s multiple comparisons was performed on the average impedance.

### 2.6 An Alternative Electrokinetic Characterization with Dielectrophoresis

For dielectrophoresis, a 3DEP analyzer (LabTech, East Sussex, UK) [29], an electrokinetic instrument similar to the potentiostat in the use of nonuniform electric fields to assess biological cells, was used to characterize the LNCaP cells. The 3DEP analyzer uses a microfluidic device with an array of 20 microwells containing 3-dimenisional circular electrodes for cell measurements. Each microwell can supply an independent frequency with the same voltage. This microfluidic device was rinsed with 100 µL of 70% ethanol (1 time), Milli-Q water (3 times), and the LCB (3 times). After which 80 µL of the resuspended LNCaP cells were introduced into the microfluidic device with a 200 µL pipet and the device was covered with an 18 mm × 18 mm glass coverslip. The microfluidic device was placed inside the 3DEP analyzer and a frequency sweep from 1 kHz to 20 MHz at 10 Vpp was completed in 60 seconds. The 3DEP analyzer outputs a relative cell response based on variations in light intensity as a function of frequency [30]. The cell response was evaluated, and outliers removed with the 3DEP analyzer software. This characterization was repeated for three biological repeats.

### 2.7 EMT Characterization with Morphology Assessment

After EMT treatment, the PC3, DU145, and LNCaP cells were rinsed with 1X DPBS then fixed with 4% paraformaldehyde solution for 10 minutes. Cells were incubated in Hoechst 33342 solution (Life Technologies, Carlsbad, CA, 1:2500 dilution in 1X DPBS) for 5 minutes to visualize the nuclei then rinsed twice with 1X DPBS. The Keyence BZ-X800 All-in-one Fluorescence Microscope was used for imaging.

### 2.8 EMT Characterization with Immunofluorescence

After EMT treatment, cells were rinsed with 1X DPBS then fixed with 4% paraformaldehyde solution for 10 minutes. Cells were rinsed in 1X DPBS then permeabilized in 0.2% Triton X for 30 minutes. The cells were washed with 1X DPBS and then incubated in blocking buffer (consisting of 5% (v/v) normal donkey serum (Jackson ImmunoResearch, West Grove, PA) and 1% (w/v) bovine serum albumin (Life Technologies, Carlsbad, CA), for 15 minutes. This was followed by incubation in primary antibodies for 1 hour at room temperature: rabbit E-cadherin polyclonal antibody (Invitrogen, Waltham, MA, 1:100 dilution), mouse ZO-1 monoclonal antibody (Invitrogen, Waltham, MA, clone A12, 1:100 dilution), rabbit N-cadherin polyclonal antibody (Invitrogen, Waltham, MA, 1:100 dilution), and rabbit vimentin polyclonal antibody (Invitrogen, Waltham, MA, 1:100 dilution), all diluted in the blocking buffer. Cells were subsequently washed 1X DPBS then incubated with secondary antibodies: donkey anti-rabbit AlexaFluor488 (Life Technologies, Carlsbad, CA), donkey anti-mouse AlexFluor594 (Life Technologies, Carlsbad, CA) diluted in 1% blocking buffer for 1 hour at room temperature. Afterwards, cells were incubated in Hoechst 33342 Solution (1:2500 dilution in 1X DPBS) for 5 minutes to visualize the nuclei then rinsed twice with 1X DPBS. The Keyence BZ-X800 All-in-one Fluorescence Microscope was used for imaging.

After imaging, the green channel intensity from the immunofluorescent images was quantified using ImageJ. Each image was imported and split into its red, green, and blue (RGB) channels. Only the green channel, corresponding to the fluorescent signal, was analyzed. The red and blue channels were discarded. To ensure consistent measurements across images, a rectangular region of interest (ROI) was defined using the rectangle tool and placed in the same location for each image. Two ROIs per image were used to assess different areas of the image and obtain an average fluorescence intensity. The average green channel intensity for each marker (E-cadherin and vimentin) was calculated and plotted to compare fluorescence levels. A one-way ANOVA with Šidák’s multiple comparisons was performed on the average intensity.

## 3. Results

The EMT was induced in PCCs and characterized using EIS (or dielectrophoresis), morphology assessments, and immunofluorescent staining; Fig. 1 depicts the EMT and outlines the experimental workflow. During EMT, cells lose epithelial features, noted by the downregulation of epithelial markers such as E-cadherin and ZO-1, and gain mesenchymal features noted by the upregulation of mesenchymal markers such as N-cadherin and vimentin. An intermediate phenotype, where both epithelial and mesenchymal markers are present, can also occur, Fig. 1.1. To initiate the EMT, PC3, DU145, and LNCaP cells were treated for 5 days with EMT-inducing media. After the EMT treatment, the cells were characterized using EIS (or dielectrophoresis) and the phenotype change was visually confirmed by nuclei stain and immunofluorescence imaging, Fig. 1.3. Supplemental Fig. S1 illustrates the method used to quantify the intensity of the fluorescent stains. Prism 10 was used to analyze the impedance spectra, the technical replicates were analyzed for outliers, normalized, and averaged.

**Figure 1.**
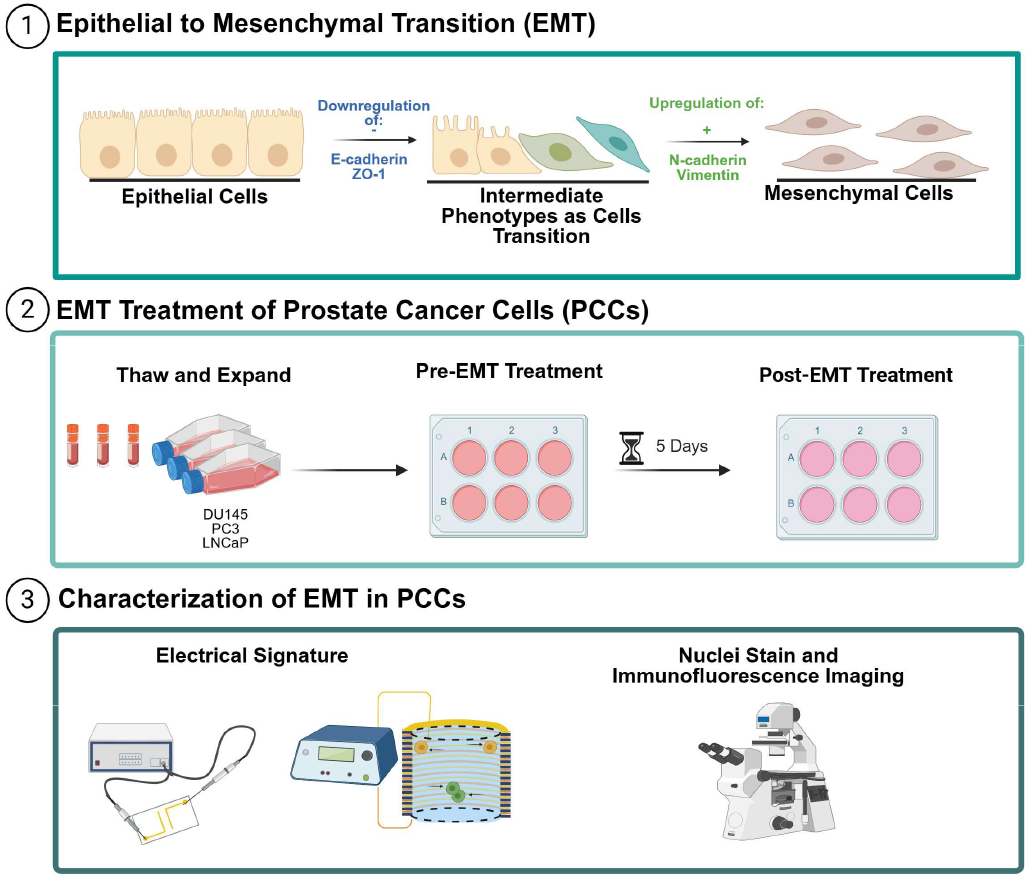
EIS experimental workflow for characterizing phenotype changes in PCCs. (1) Illustration of EMT in cancer cells. Initially, cells exhibit an epithelial phenotype characterized by the expression of E-cadherin and ZO-1. Then, the cells undergo downregulation of E-cadherin and ZO-1, transitioning to an intermediate phenotype. In this stage there is a shift in the protein expression profile with an upregulation of N-cadherin and vimentin leading to mesenchymal phenotype. (2) DU145, PC3, and LNCaP cells were obtained from cryogenic storage, thawed, and expanded in proliferation media. Cells were seeded with EMT-inducing media and allowed 5 days to incubate. (3) Cells were characterized using EIS and the 3DEP analyzer. A phenotype change was validated by a nuclei stain and immunofluorescence imaging. Figure created with Biorender.com.

### Electrical Signature of PCCs

The impedance spectra of PCCs were collected, normalized to remove dependency on LCB, and averaged. Fig. 2 shows the unnormalized and normalized Bode plot over the frequency range 350 Hz – 5 MHz of one passage (n=1 biological replicate) of PC3 EMT+ and DU145 EMT+ cells. For both PC3 and DU145 cells, the impedance decreases with increased frequency, displaying the typical s-shape impedance spectrum. Comparing Fig. 2A and 2C, the impedance spectra of the PC3 and DU145 cells are defined by two transitions (102-103 Hz and 106-107 Hz) and a plateau (103-106 Hz). This trend remains when comparing the normalized plots, Fig. 2B and 2D. The technical replicates for the PC3 cells have a smaller spread than DU145 cells.

**Figure 2.**
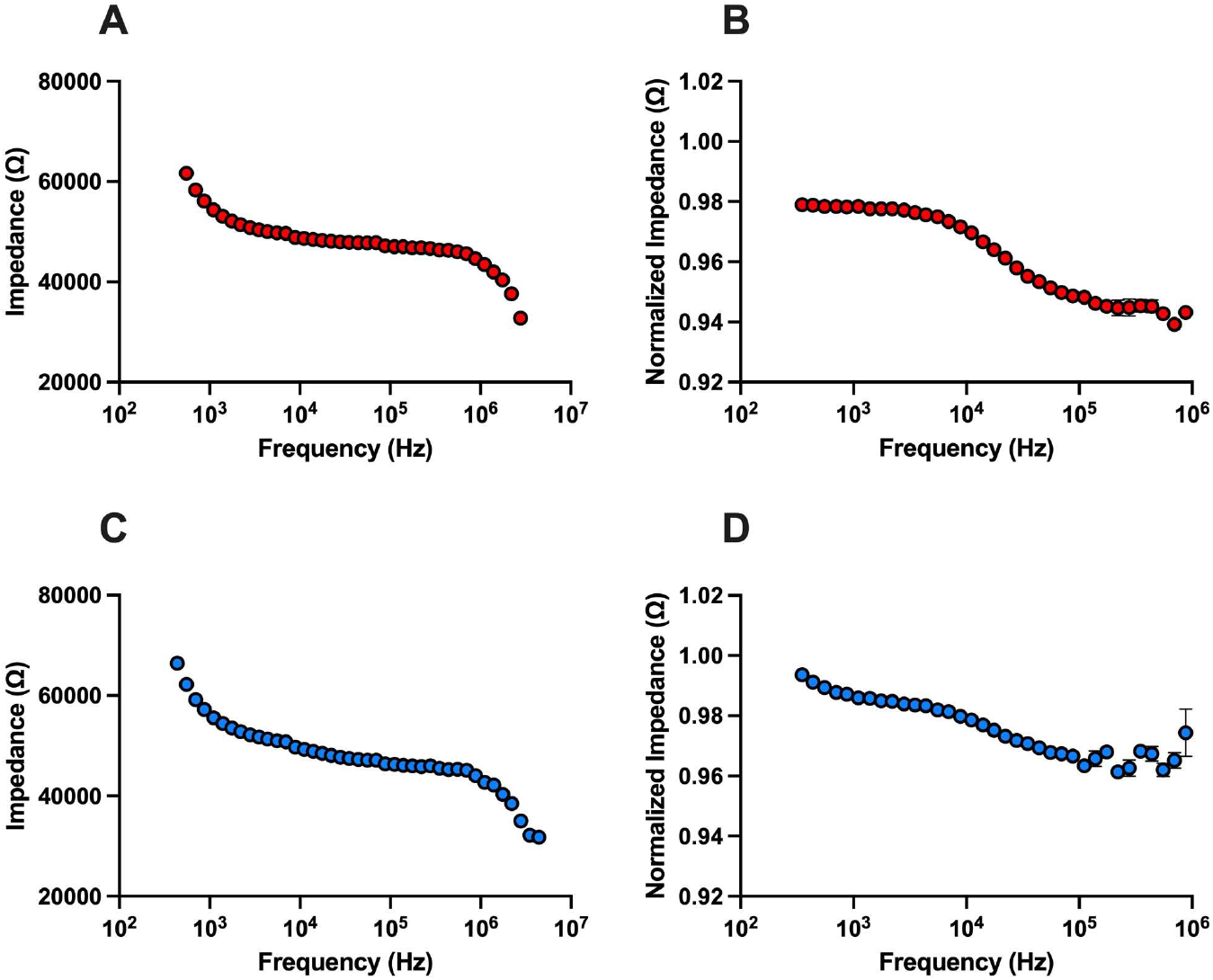
EIS cell analysis of PCCs. (A) Unnormalized and (B) normalized EIS spectrum of PC3 EMT+ cells (n=1). (C) Unnormalized and (D) normalized EIS spectrum of DU145 EMT+ cells (n=1). In all plots, data points represent average impedance, technical replicates equal 3 to 5 individual measurements and error bars are standard error mean. Most error bars in (A-D) are too small to be visualized.

We completed the EMT treatment and measured the impedance signature of PC3 and DU145 cells. Fig. 3A and 3B present the normalized impedance of three biological replicates (n=3) of the PC3 and DU145 cells, respectively. EMT+ signifies cells were treated with EMT induction media and EMT-signifies cells were not treated. Similar to Fig. 2, the impedance decreased with increased frequency. There is a small separation in the impedance spectra for the PC3 EMT+ and PC3 EMT-cells with the error bars overlapping. In contrast, there is a noticeable separation in the impedance spectra for the DU145 EMT+ and DU145 EMT-cells and the error bars do not overlap. Fig 3C and 3D show the impedance spectra of PC3 and DU145 cells compared to the epithelial (HPrECs) and mesenchymal (hMSCs) controls. HPrECs serve as the negative control for EMT induction while hMSCs serve as the positive control for EMT induction. The impedance spectrum of the mesenchymal control is higher than that of the epithelial control. The averaged impedance spectrum of DU145 EMT-cells falls below and overlaps the epithelial control. The DU145 EMT+ cells initially overlap with the epithelial control but shifts upward with increasing frequency, moving closer to the mesenchymal control. The error bars of the DU145 EMT-cells overlap with the error bars of the epithelial control. In contrast, the average impedance spectra of PC3 EMT-cells and PC3 EMT+ cells overlap the mesenchymal control. The error bars of the PC3 EMT-cells and PC3 EMT+ cells overlap with each other and the mesenchymal control. In Fig. 3E and 3F, the averaged normalized impedance of PC3 EMT- and PC3 EMT+ cells are similar, while the average normalized impedance of DU145 EMT- and DU145 EMT+ cells differ with statistical significance (^**^ p < 0.05). Also, the average normalized impedance for the epithelial and mesenchymal control are different with statistical significance. Each DU145 condition (EMT- and EMT+) differ significantly from both the epithelial and mesenchymal controls.

**Figure 3.**
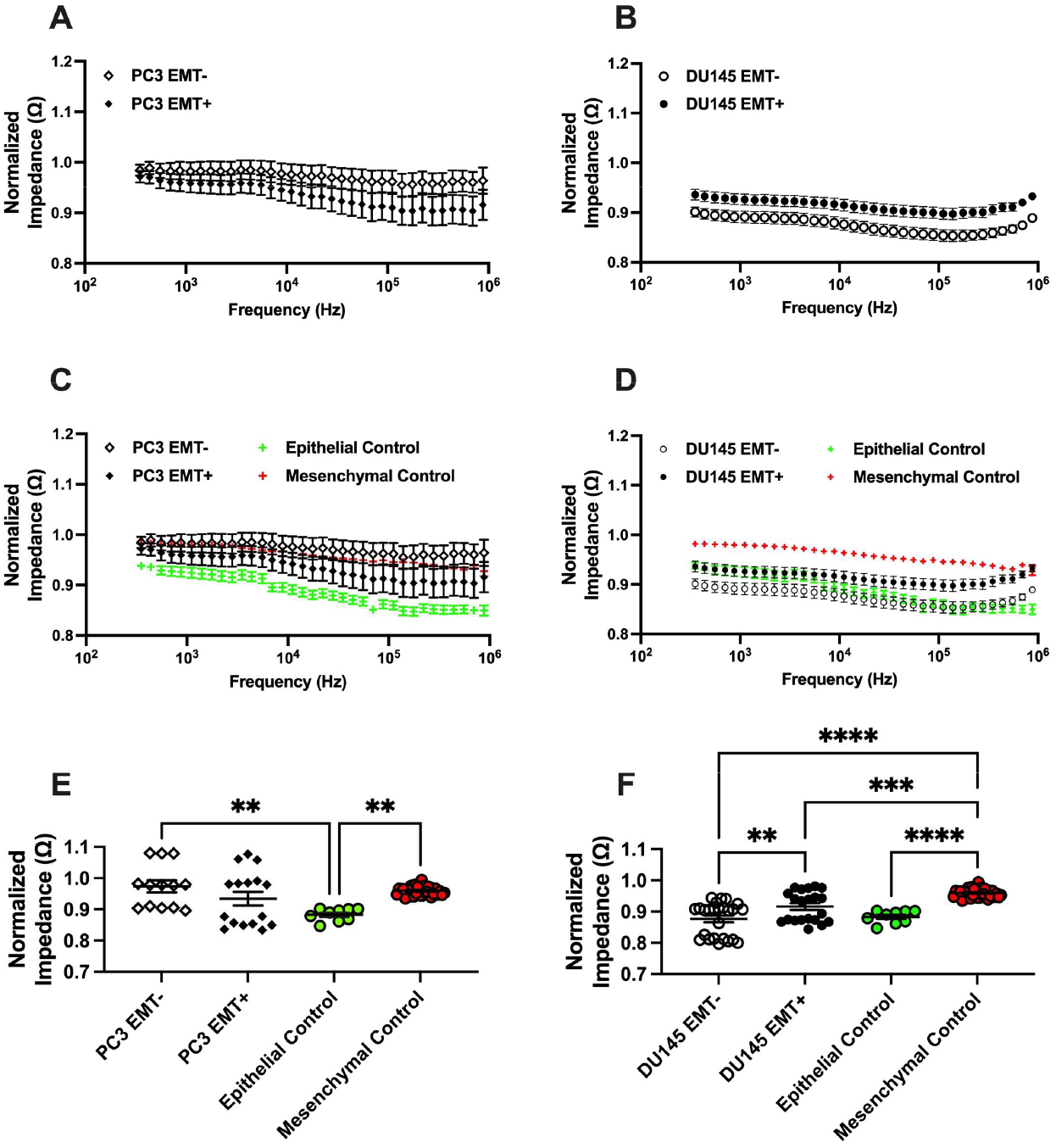
Normalized EIS cell analysis of EMT treated PCCs. (A) Average spectra of PC3 EMT-cells and PC3 EMT+ cells. (B) Average spectra of DU145 EMT-cells and DU145 EMT+ cells. Average spectra of (C) PC3 EMT- and PC3 EMT+ cells and (D) DU145 EMT-and DU145 EMT+ cells compared to epithelial and mesenchymal controls. Average normalized impedance of (E) PC3 EMT-and PC3+ cells and (F) DU145 EMT-and DU145 EMT+ cells compared to epithelial and mesenchymal controls. Error bars represent standard error mean. n=3 for PC3, DU145, epithelial control, and mesenchymal control cells. Statistical analysis completed on pooled data sets; ^**^ p < 0.05, ^***^ p < 0.001, and ^****^ p < 0.0001.

The electrical signature of LNCaP cells treated with EMT induction media was determined with EIS and the 3DEP analyzer. Small differences at select frequencies were observed for both the impedance and DEP spectra. At higher frequencies, the impedance spectra show that LNCaP EMT-cells have higher impedance than LNCaP EMT+ cells, with overlap occurring at lower frequencies, Supplemental Fig. S2. In the DEP spectra, LNCaP EMT-cells exhibit higher light intensity response at certain frequencies, while at lower frequencies LNCaP EMT+ cells show a greater response, Supplemental Fig. S3.

To further assess the electrical signature, additional EIS measurements were performed on mixtures of PC3 EMT-/+, DU145 EMT-/+, and LNCaP EMT-/+ cells. Supplemental Fig. S4 shows that differences in the impedance spectra are observed at higher frequencies for DU145 EMT-, DU145 EMT+, and DU145 EMT-/+ cells, as well as for LNCaP EMT-, LNCaP EMT+, and LNCaP EMT-/+ cells. At lower frequencies, there is overlap among the three conditions for both DU145 and LNCaP cells. For PC3 cells, the impedance spectra of EMT-/+ mixtures align closely with those of EMT-cells.

### Biological Assessment of EMT Treatment

It is necessary to characterize the EMT using biological markers. Thus, we examined the morphology and protein expression of the PCCs after EMT treatment. Fig. 4 presents nuclei stains of PC3, DU145, and LNCaP cells. The PC3 EMT- and EMT+ cells do not exhibit any differences in their morphology, they are spindle-like, fibroblastic. For the DU145 cells, the EMT-cells have a cobblestone morphology while EMT+ cells feature a spindle-like, fibroblastic morphology. The LNCaP EMT-cells show a similar cobblestone morphology, while the LNCaP EMT+ cells show a more fibroblastic morphology.

**Figure 4.**
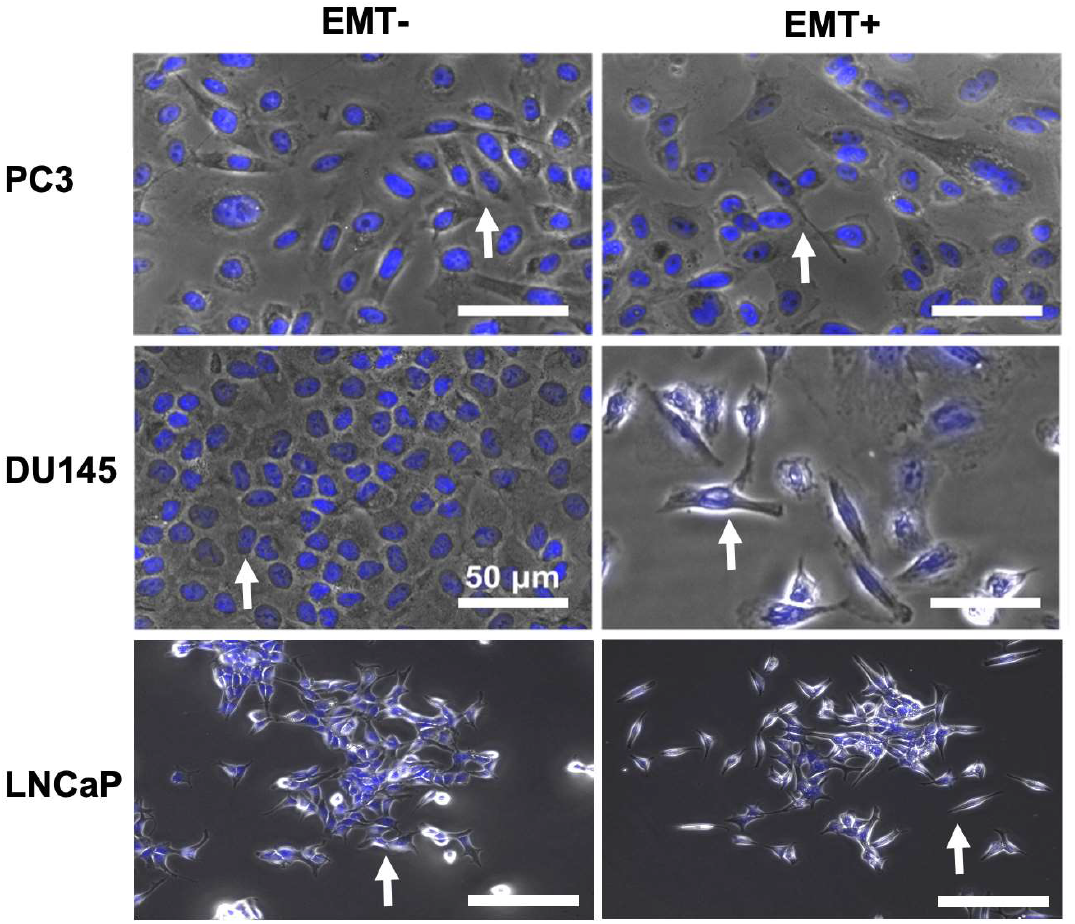
Morphology assessment of EMT treated PCCs. Phase contrast images overlayed with Hoechst-stained nuclei of PC3, DU145, and LNCaP cells without (EMT-) and with EMT (EMT+) treatment. The white arrows indicate a representative cell exhibiting characteristic epithelial morphology under EMT-conditions and mesenchymal morphology under EMT+ conditions.

Fig. 5 displays the protein expression of E-cadherin and vimentin for the PCCs with immunofluorescence staining and quantification. For the PC3 cells, a decrease in fluorescence is observed and quantified for both E-Cadherin and vimentin. The DU145 cells show positive fluorescence for E-cadherin and vimentin; however, the E-cadherin expression decreased following EMT treatment, while vimentin expression increased. Similarly, LNCaP cells exhibited positive fluorescence for E-cadherin and vimentin, with a decrease in E-cadherin after EMT treatment and increased vimentin expression. ZO-1 was used as an additional epithelial marker, and downregulation was observed in DU145 EMT+ cells, Supplemental Fig. S5. N-cadherin was used as an additional mesenchymal marker, and upregulation was observed in LNCaP EMT+ cells, Supplemental Fig. S6.

**Figure 5.**
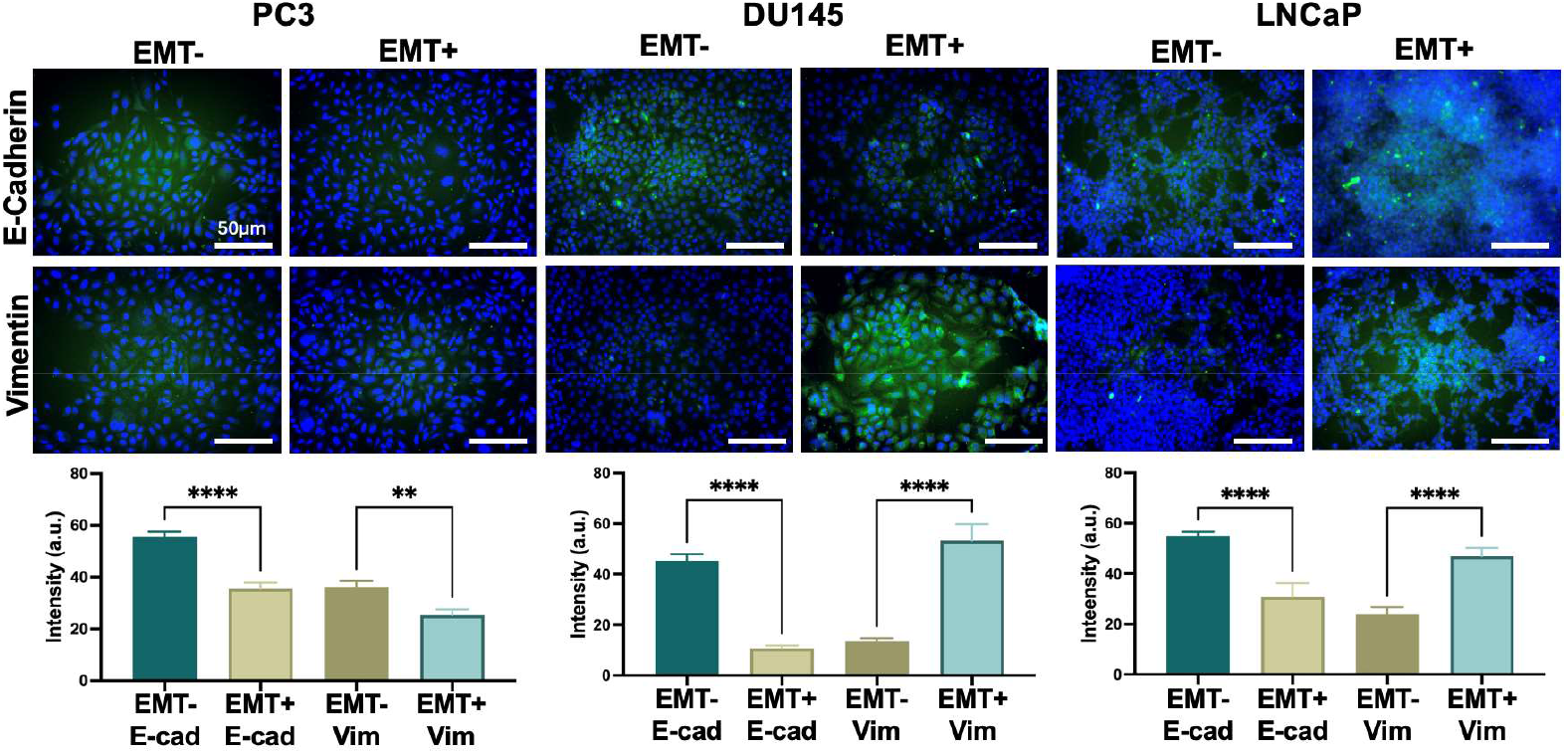
Immunofluorescent staining of PCCs without and with EMT treatment (EMT- and EMT+, respectively). The staining highlights the expression of epithelial marker E-cadherin and mesenchymal marker vimentin both tagged with fluorescent labels. Quantification of fluorescent intensity for each marker is provided in the bar charts. Error bars represent standard error mean. n=3 for all conditions. Statistical analysis completed on pooled data sets; ^**^ p < 0.05, and ^****^ p < 0.0001.

## 4. Discussion

Characterizing the electrical signature of phenotype changes, such as the EMT, in PCCs is essential as it correlates with chemoresistance [6]. The EMT involves the transition of cells through multiple states marked by morphological, protein, gene, and electrical changes in their phenotype. EIS effectively analyzes cell phenotypes by measuring impedance induced by internal and external factors. To capture changes in the electrical signature associated with EMT, we examined frequencies from 350 Hz to 5 MHz. This range detects changes in the cell membrane properties potentially related to the loosening of tight junctions and changes in membrane structure. We chose PC3, DU145, and LNCaP cell lines based on their varying metastatic potentials [25, 31, 32] and levels of cancer stemlike cells [33]. Also, these cells are frequently used in PCa research [34] making them suitable for this study. Thus, this study focused on inducing the EMT in PC3, DU145, and LNCaP cells and detecting a phenotype change with EIS or dielectrophoresis. The novelty of this study is based on the label-free engineering measurement of cells’ electrical signature combined with morphology and immunofluorescent assessments to detect PCC phenotype changes related to EMT.

In this study, we treated PCCs with EMT induction media and identified cell phenotype changes using EIS. Our results show that the impedance of PC3 EMT+ cells is higher than the DU145 EMT+ cells. Similarly, we observed that the impedance of PC3 EMT-cells is higher than the DU145 EMT-cells. These findings represent Day 5 EIS measurements. In contrast, our previous study, which included Day 1 and Day 7 EIS measurements with no EMT treatment, showed that the impedance of DU145 cells was higher than PC3 cells on Day 1 [9], a finding also supported by literature [19]. By Day 7, the impedance spectra indicated a higher impedance for PC3 cells compared to DU145 cells [9]. This shift aligns with our current findings and underscores the impact of cell culture duration on EIS measurements, confirming the consistency of impedance trends across studies. Additionally, it should be noted that EIS is sensitive enough to detect subtle day-to-day changes in cells.

EIS distinguished EMT+ and EMT-cells for the DU145 cells (Figure 3B, 3D, and 3F) indicating a phenotype change. Similarly, the electrical signature of LNCaP cells, determined with EIS and dielectrophoresis, distinguished EMT+ and EMT-cells (Supplemental Fig. S2 and S3). However, the EIS of EMT+ and EMT-cells for PC3 cells was inclusive. Additional comparisons to epithelial and mesenchymal controls provided more evidence of a phenotype change in the DU145 cells. Supplemental Figure S4 provides additional evidence of EIS distinguishing EMT+ and EMT-cells through the analysis of mixtures of DU145 and LNCaP cells. As seen in Fig. 4, there are morphological differences between the DU145 EMT-cells and the LNCaP EMT-cells featuring cobblestone morphology while DU145 EMT+ cells and LNCaP EMT+ cells display spindle-like, fibroblastic morphology. These alterations in morphology were due to the downregulation of E-cadherin in both DU145 and LNCaP cells (Fig. 5), ZO-1 in DU145 cells (Supplemental Fig. S5), and the upregulation in N-cadherin in LNCaP cells (Supplement Fig. S6). Thus, we successfully induced and detected the occurrence of EMT in DU145 and LNCaP cells, distinguishing between epithelial state cells (EMT-) and mesenchymal state cells (EMT+). There was no phenotype change induced in PC3 cells because they were already in the mesenchymal state. To our knowledge, this is the first study to utilize EIS to detect EMT in PCCs, demonstrating selectivity in distinguishing between epithelial (EMT-), mesenchymal (EMT+), and mixture (EMT-/+) phenotypes in DU145 and LNCaP cells. These results align with hallmark impedimetric literature concerning phenotype changes during stem cell maturation [35-37].

Previous studies utilizing impedimetric techniques have demonstrated their utility in discerning EMT states in various cancer models [20, 21], underscoring the importance of electrical signatures in understanding cellular phenotypic plasticity. Specifically, Schneider et al. [20] treated lung (A549) and breast (MDA-MB231) cancer cells with TGF-β to induce EMT and characterized phenotype changes with morphology assessment, F-actin and ZO-1 protein expression, and an ECIS device with an array of circular electrodes. The A549 cells exhibited a shift from a cobblestone to a fibroblastic morphology, increased F-actin fibers, and decreased ZO-1 protein expression upon TGF-β treatment. ECIS enabled real-time, time-resolved impedance measurements of the adherent cells’ dynamic responses to TGF-β, confirming the induction of EMT in A549 cells while no changes were noted in the more aggressive MDA-MB231 line. Similarly, Zhao et al. [21] used IFC to detect EMT in A549 cells induced with TGF-β, discerning differences between A549 and A549 EMT based on membrane capacitance and cytoplasm conductivity. The impedance differences observed by Zhao et al. were validated through microscopy, providing an analysis of EMT-induced changes in electrical properties.

Our work extends these investigations by applying EIS and dielectrophoresis to PCCs, offering a steady-state analysis of EMT-specific electrical changes. While Schneider et al. focused on the dynamic, real-time impedance changes, our approach provides a complementary perspective by evaluating steady-state electrical signatures of EMT- and EMT+ states in PCCs. Additionally, the use of parallel electrode microfluidic device in our study allows for label-free analysis reducing the need for antibody labeling, streamlining sample preparation and maintaining cell integrity. This contrasts with Zhao et al.’s single-cell IFC measurements, as our EIS and dielectrophoresis measurements provide an alternative platform for identifying EMT in bulk PCC populations without need for fluid flow. Furthermore, our study confirmed the induction of EMT in PCCs through typical morphology changes—a shift from a cobblestone to spindle-like phenotype—as well as through protein expression markers, including downregulation of E-cadherin and upregulation of vimentin. These phenotypic shifts align with typical EMT-associated changes reported in other cancer models [22, 38, 39], underscoring the relevance of our findings. By comparing our steady-state EIS data to the dynamic and single-cell measurements of Schneider et al. and Zhao et al., our work broadens the scope of electrokinetic techniques to study EMT across different cancer types.

In the context of these previous studies, our research reinforces the utility of EIS in distinguishing between EMT+ and EMT- phenotypes in cancer cells, contributing to a rapidly growing body of literature on the application of label-free technologies in oncological research. Despite the advancements demonstrated by impedimetric techniques in characterizing EMT, challenges such as standardizing measurement protocols and interpreting complex impedance spectra persist. Our study’s limitations, including variability in cell line responses and the need for larger sample sizes (such as fresh patient samples), emphasizing the importance of addressing these constraints to fully leverage impedance spectroscopy in elucidating the dynamics of EMT and its implications for chemoresistance and cancer recurrence in PCa. Translating these insights into a clinical setting may require additional improvements in the EIS technology we use to ensure adequate sample preparation and developing cancer cell monitoring devices. However, there is value in the simplicity of our simple parallel electrode device.

Despite these limitations, our findings have clinical relevance. EIS can be utilized to monitor and detect cancer cell dynamics in real time. Information from these studies can aid in the investigation of the molecular dynamics of EMT and provide more insight to when in time this program occurs in cancer cells. Characterizing the electrical signature of EMT can aid in creating drugs that target cancer cells susceptible to EMT; this strategy is underway for other cancers [7]. EIS can also be employed for drug development by assessing how effectively drugs target cancer cells undergoing EMT. This approach aligns with pharmaceutical industries’ interest in developing drugs targeting specific cancer cell phenotypes. EIS extends to various clinical applications, including assessing EMT-associated metastatic potential through cell detachment and facilitating liquid biopsies. Coupling EIS with techniques like dielectrophoresis enables label-free processing and monitoring of cancer cells in patient samples, offering new insights into phenotypic characterization such as visualization of heterogeneity.

## 5. Conclusions

EIS represents a promising tool for detecting and monitoring EMT in PCCs. This study presents important advancements, including the application of electrokinetic techniques (EIS and dielectrophoresis) for label-free EMT detection, characterization of electrical signatures of PCCs following EMT, and demonstrated sensitivity of these approaches to cellular transitions. Integrating impedimetric assays with traditional cancer diagnostic methods may enhance early detection of cancer recurrence and enable timely, EMT-targeted interventions, which are crucial for the long-term care of PCa patients. Future investigations will focus on employing label-free microfluidic cell sorting strategies to selectively isolate EMT-sensitive cells for further characterization. This will include advanced electrokinetic analysis (generation of impedance and dielectrophoresis spectra), molecular profiling (protein and gene expression of epithelial and mesenchymal markers), and imaging techniques (scanning electron microscopy). The advanced electrokinetic analysis will include microfluidic devices with single-cell and bulk cell modalities, increased throughput, and increased sensitivity of the sensing electrodes to better identify EMT in PCCs.

## Author Contributions

Conceptualization, L.L.C., L.A.H., M.T., T.N.G.A; methodology, L.L.C., L.A.H., M.T., T.N.G.A.; formal analysis, L.L.C., L.A.H., M.T; investigation, L.L.C., L.A.H., M.T.; resources, T.N.G.A.; data curation, L.L.C., L.A.H., M.T.; writing—original draft preparation, L.L.C., L.A.H., M.T., T.N.G.A.; writing—review and editing, L.L.C., L.A.H., M.T., T.N.G.A.; supervision, T.N.G.A; funding acquisition, T.N.G.A. All authors have read and agreed to the published version of the manuscript.

## Acknowledgments

The authors would like to thank UCI’s Integrated Nanosystems Research Facility for support with microfluidic device fabrication.

