## Supplemental Information for "Electrical Impedance Spectroscopy as a Tool to Detect the Epithelial to Mesenchymal Transition in Prostate Cancer Cells"

The workflow for quantifying the fluorescence intensity of the immunofluorescent stains is illustrated. Images are imported into ImageJ, split into their red, green, and blue channels, and analyzed using only the green channel. ROIs are drawn to measure the fluorescence intensity, and the mean intensity is calculated for more analysis.

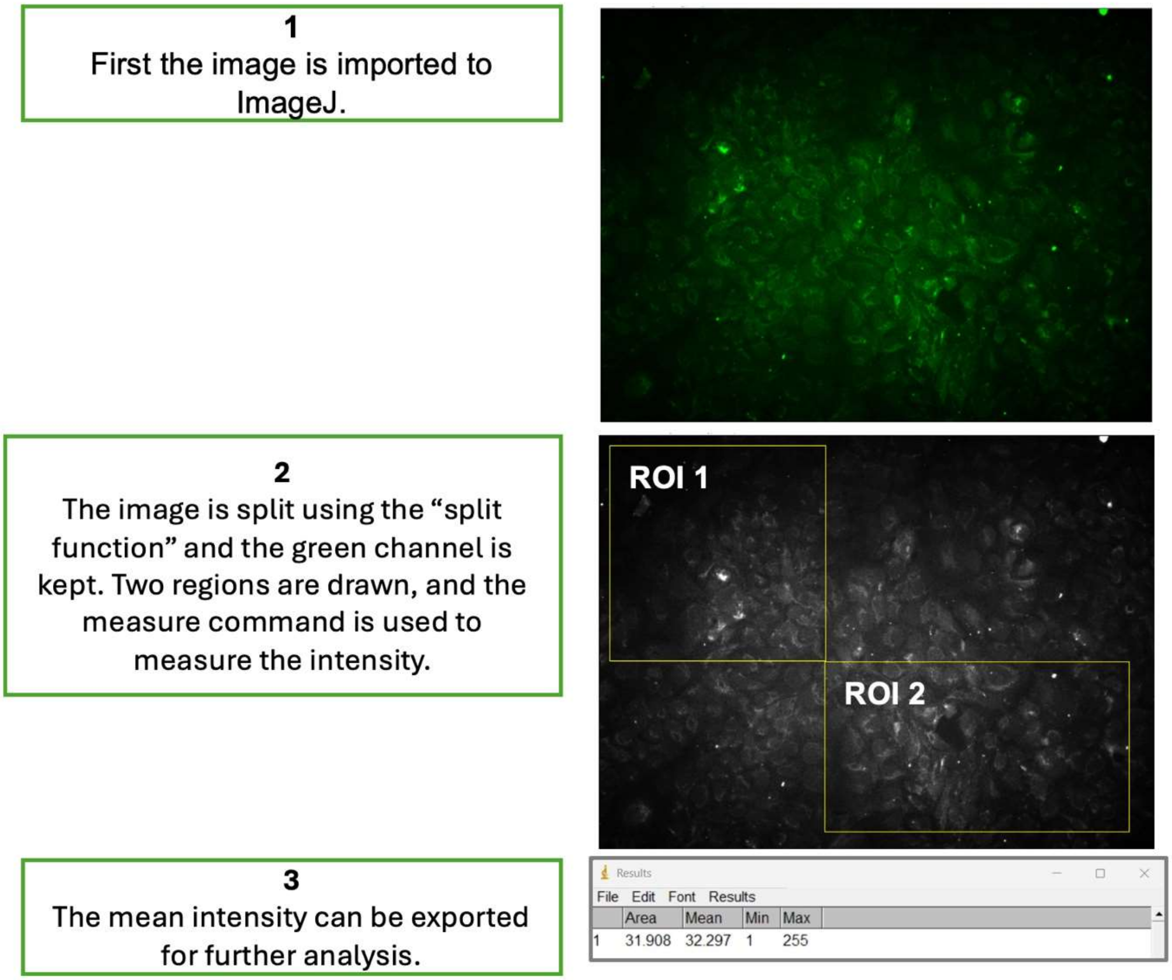


**Figure S1.** Workflow for quantifying the fluorescence intensity of immunofluorescent stains.

EIS and the 3DEP analyzer were used to determine the electrical signature of LNCaP cells without and with EMT treatment. Subtle differences in the electrical signature were observed at specific frequencies, denoted with black dashed line boxes, for both the DEP and impedance spectra.

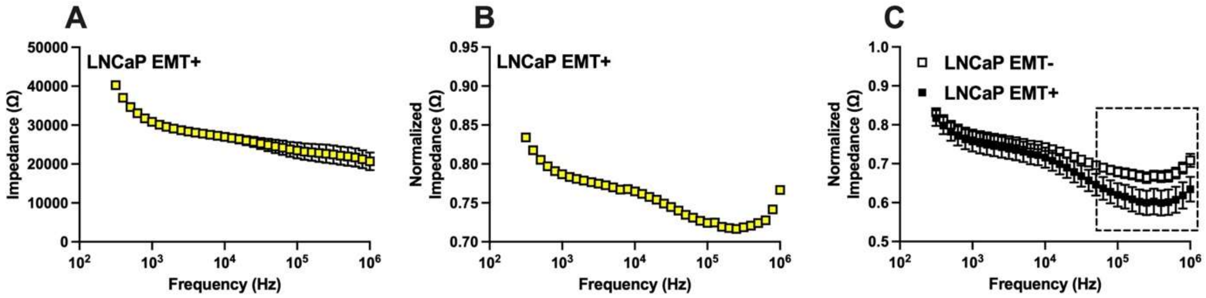


**Figure S2.** EIS analysis of EMT treated LNCaP cells. (A) Unnormalized and (B) normalized EIS spectrum of LNCaP EMT+ cells (n=1). (C) Average spectra of LNCaP EMT- cells and LNCaP EMT+ cells.

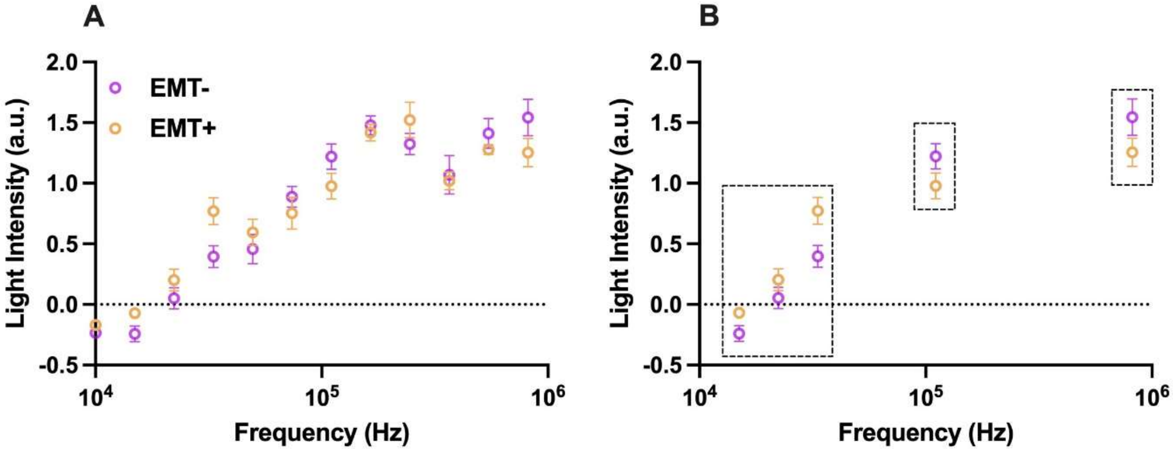


**Figure S3.** 3DEP analysis of EMT treated LNCaP cells. (A) Average light intensity response spectra of LNCaP EMT- and EMT+ cells. (B) Black boxes highlight specific frequencies that emphasize differences in the electrical signature between EMT- and EMT+ conditions.

EIS analysis was completed on mixtures of PC3 EMT-/+, DU145 EMT-/+, and LNCaP EMT-/+ cells. Differences are observable at higher frequencies for DU145 and LNCaP cells across EMT-, EMT+, and EMT-/+ conditions. At lower frequencies there is considerable overlap among these conditions. In the case of PC3 cells, the impedance of EMT-/+ mixture is similar to those of EMT- cells. The buffer is included for reference.

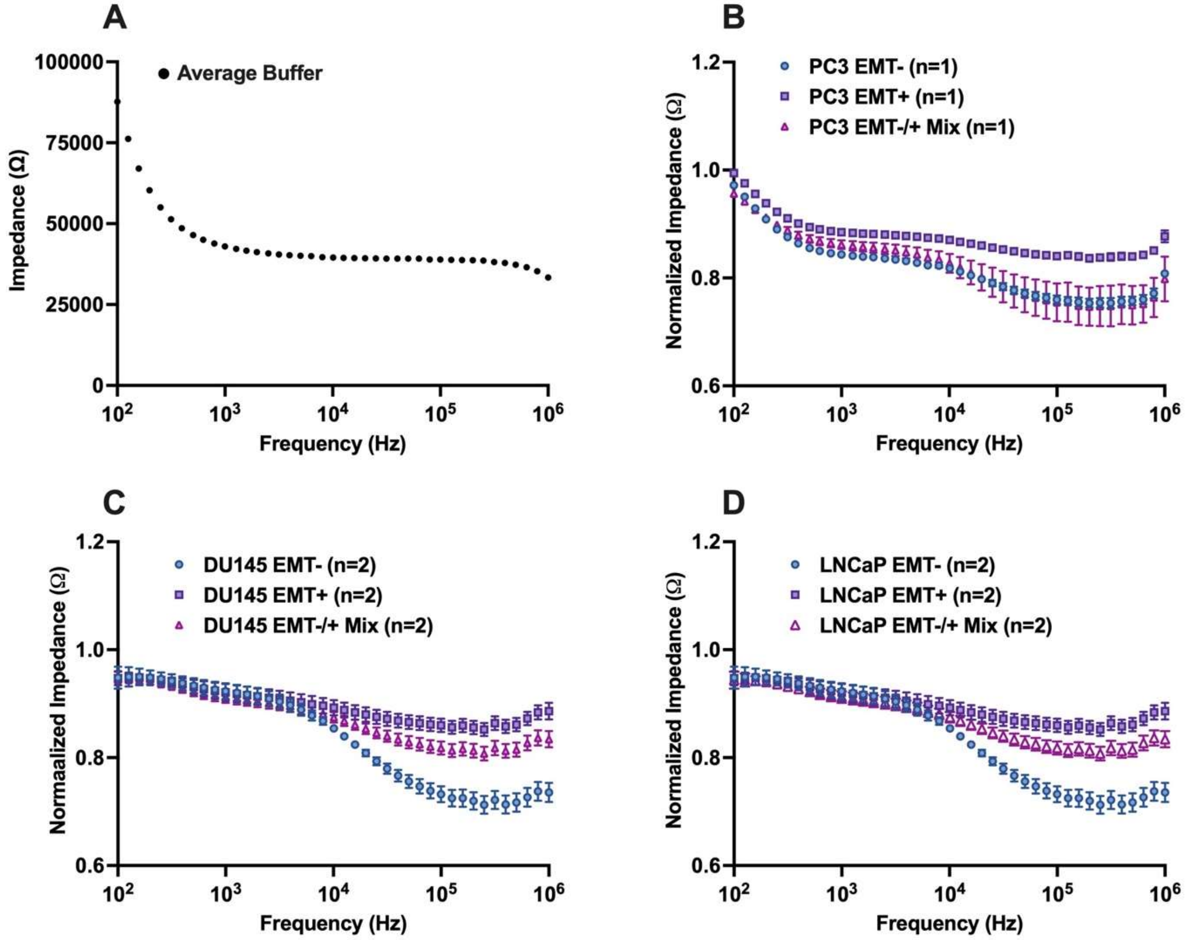


**Figure S4.** EIS analysis of mixtures of EMT- and EMT+ cells. (A) Impedance spectra of average buffer. Normalized impedance spectra of mixtures for (B) PC3 (n=1), (C) DU145 (n=2), and (D) LNCaP cells (n=2).

For the DU145 cells, ZO-1 was used as an additional indicator of EMT. A reduction in ZO-1 protein expression was observed following EMT treatment indicating that the DU145 cells transitioned from an epithelial phenotype to a mesenchymal phenotype.

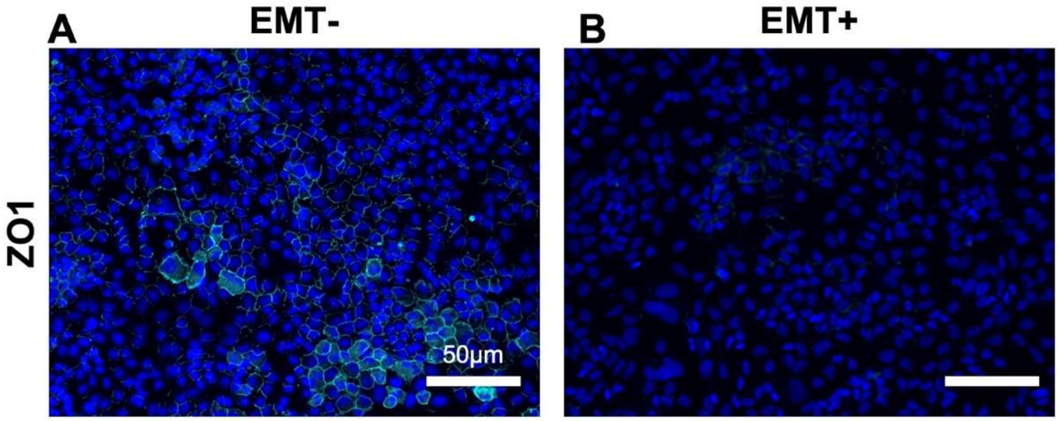

**Figure S5.** Immunofluorescent staining of DU145 cells without and with EMT treatment (EMT- and EMT+, respectively). The staining highlights the protein expression of the epithelial marker ZO-1.

For the LNCaP cells, N-cadherin served as another indicator of EMT. An increase in N-cadherin protein expression was observed with EMT treatment suggesting that the LNCaP cells underwent a shift from an epithelial phenotype to a mesenchymal phenotype.

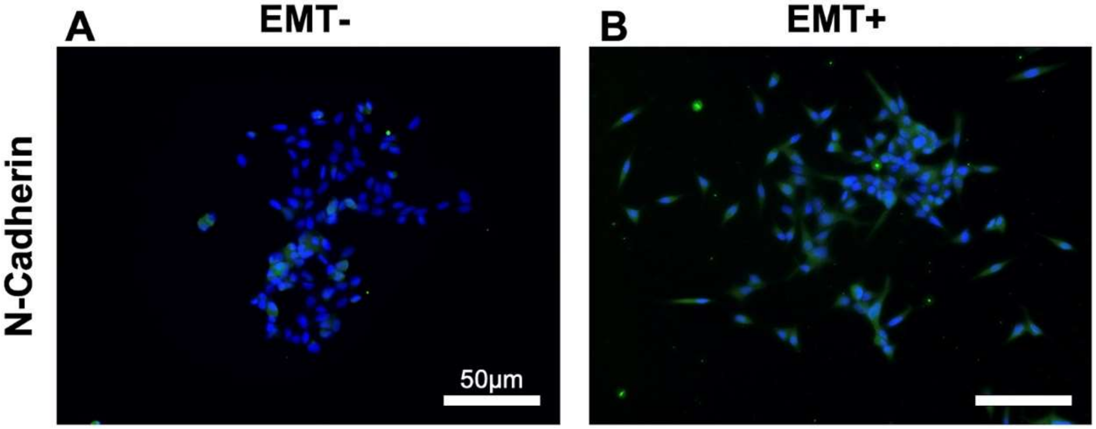


**Figure S6.** Immunofluorescent staining of LNCaP cells without and with EMT treatment (EMT- and EMT+, respectively). The staining highlights the expression of the mesenchymal marker N-cadherin.
